# L-DOPA induces spatially discrete changes in gene expression in the forebrain of mice with a progressive loss of dopaminergic neurons

**DOI:** 10.1101/2025.02.13.638140

**Authors:** Anna Radlicka-Borysewska, Magdalena Ziemiańska, Mateusz Zięba, Łukasz Szumiec, Monika Bagińska, Magdalena Chrószcz, Sławomir Gołda, Jacek Hajto, Michał Korostyński, Grzegorz Kreiner, Joanna Pera, Marcin Piechota, Jan Rodriguez Parkitna

## Abstract

L-3,4-Dihydroxyphenylalanine (L-DOPA) is effective at alleviating motor impairments in Parkinson’s disease (PD) patients but has mixed effects on nonmotor symptoms and causes adverse effects after prolonged treatment. Here, we analyzed the spatial profile of L-DOPA-induced gene expression in the forebrain of mice with an inducible progressive loss of dopaminergic neurons (the TIF-IA^DATCreERT2^ strain), with a focus on the similarities and differences in areas relevant to different PD symptoms. The animals received a 14-day L-DOPA treatment, and 1 h after the final drug injection, a spatial transcriptome analysis was performed on coronal forebrain sections. A total of 121 genes were identified as being regulated by L-DOPA. We found that the treatment had widespread effects extending beyond the primary areas involved in dopamine-dependent movement control. An unsupervised clustering analysis of the transcripts recapitulated the forebrain anatomy and indicated both ubiquitous and region-specific effects on transcription. The changes were most pronounced in layers 2/3 and 5 of the dorsal cortex and the dorsal striatum, where a robust increase in the abundance of activity-regulated transcripts, including *Fos*, *Egr1*, and *Junb*, was observed. Conversely, transcripts with a decreased abundance, e.g., *Plekhm2* or *Pgs1*, were identified primarily in the piriform cortex, the adjacent endopiriform nucleus, and the claustrum. Taken together, our spatial analysis of L-DOPA-induced alterations in gene expression reveals the anatomical complexity of treatment effects, identifying novel genes affected by the drug, as well as molecular activation in brain areas relevant to the nonmotor symptoms of PD.

## INTRODUCTION

Parkinson’s disease (PD) is characterized by the progressive degeneration of neurons in the central and autonomous nervous systems, affecting primarily midbrain dopaminergic cells but also serotonergic, noradrenergic and cholinergic neurons [1–4]. The development of PD is associated with a variety of motor and nonmotor symptoms [5–7]. The cardinal motor symptoms of PD, namely, muscle rigidity, bradykinesia, and resting tremor, are attributed primarily to the loss of dopaminergic innervation originating in the ventral midbrain. The nonmotor symptoms of PD may precede the motor impairment and include attention and cognitive deficits, anxiety, depression, apathy, loss of smell, sleep disorders, and constipation. The etiology of nonmotor symptoms is not well understood, and the role of dopaminergic neuron degeneration in their development is often unclear.

The treatment of PD patients relies on enhancing dopaminergic neurotransmission with L-DOPA, a dopamine precursor [7–9]. L-DOPA is initially highly effective at reducing motor impairments; however, after prolonged treatment and with the progressing loss of dopaminergic neurons, the efficacy of L-DOPA decreases and adverse effects begin to develop, notably dyskinesias [10,11]. The mechanism of action of L-DOPA is more complex than could be intuitively inferred from its ability to act as a dopamine precursor, as it involves noncanonical uptake and metabolism pathways, while PD simultaneously induces changes in dopamine receptor sensitivity [12,13]. Accordingly, the mechanism of L-DOPA action in different brain areas depends on the abundance and types of monoamine transporters, as well as the distribution of D1- or D2-like receptors [14]. This finding is particularly relevant to the effects of L-DOPA on nonmotor symptoms, which are often related to the effects of PD outside the nigrostriatal pathway and likely involve the mesolimbic and corticolimbic pathways, discrete areas of the cortex, and monoamine signaling in multiple forebrain areas, including the dorsal cortex [15]. Thus, while the benefits of L-DOPA treatment are attributed to restoring balanced signaling in the nigrostriatal pathway, its effects on the signaling involved in nonmotor symptoms are complex and differ depending on the pathway affected.

An analysis of the effects of L-DOPA on neuronal gene expression is essential for elucidating its mechanism of action. While treatment with dopamine receptor agonists has relatively minor effects on gene expression in the striatum with intact dopaminergic innervation [16,17], studies of dopamine-depleted animal models have shown robust changes in the transcript profile in the striatal and cortical areas after dopaminergic treatment. After chemical or genetic lesioning of dopaminergic neurons, acute L-DOPA treatment strongly induces activity-dependent gene expression (i.e., immediate early genes, including *Fos* or *Egr1*), primarily through the sensitization of D1 receptors in the nigrostriatal pathway [18–21]. These effects are most pronounced in the dorsolateral striatum, which controls the execution of movement sequences, and thus is consistent with the drug’s efficacy in alleviating motor symptoms. L-DOPA-induced gene expression in dopamine-depleted mice has also been reported in other brain areas, including the prefrontal cortex, motor cortex, globus pallidus, subthalamic nucleus, amygdala and piriform cortex [18,19,22,23]. Studies using array- and RNA-seq-based gene expression profiling confirmed the extensive activation of activity-regulated transcripts in the striatum, e.g., [24–31], and, to an extent, in the prefrontal cortex [32]. Together, these studies reveal a complex pattern of differentially expressed transcripts that varies depending on the model organism, specific subareas selected for analysis, method of dopaminergic lesion induction and L-DOPA treatment regimen. Two studies are particularly notable for assessing the effects of L-DOPA on gene expression in specific cell types, either by the use of translational ribosome affinity purification [29] or single-nucleus RNA sequencing [31]. Both reports show strong effects of L-DOPA on neuronal cells expressing the dopamine D1 receptor in the dorsal striatum (i.e., putative medium spiny neurons of the direct pathway) and focus on the correlation between the expression patterns and the development of dyskinesias. The analysis of single nuclei in particular clearly revealed a nonhomogeneous response to L-DOPA even within the population of D1-expressing neurons, with specific neuron subpopulations showing a particular persistence of changes in gene expression (i.e., expressing *Act* and markers of the striatal matrix or patches). Nevertheless, a limitation of these studies is that they focus on only a single brain area affected by PD and L-DOPA treatment, and even within the targeted area, the spatial pattern of the observed changes remains uncertain. Moreover, the two highlighted studies, as well as the great majority of transcriptome profiling studies, examine the striatum. This choice is justified by the importance of the striatum in the etiology of PD symptoms and the mechanism of L-DOPA action; however, reports based on in situ hybridization and immunohistochemical techniques have shown that changes in gene expression are also observed in other frontal brain structures.

In this study, we used spatial transcriptomics to determine the effects of chronic L-DOPA treatment on the forebrain transcriptome in the TIF-IA^DATCreERT2^ mouse model with a tamoxifen-induced progressive loss of dopaminergic neurons in the midbrain. We find that drug-induced changes in gene expression in the forebrain include both ubiquitous and subregion-specific patterns, confirming some observations and revealing previously overlooked targets. Thus, we establish a spatial fingerprint of L-DOPA action on the PD-like forebrain, showing distinct effects on forebrain areas with potential relevance to nonmotor symptoms.

## MATERIALS AND METHODS

### Animals

TIF-IA^DATCreERT2^ male mice were housed at the animal facility of the Maj Institute of Pharmacology PAS under the following conditions: an air temperature of 22±2 °C, an air humidity of 55±10%, and a 12/12 h light/dark cycle. TIF-IA^DATCreERT2^ mice were bred on a C57BL/6N background as described previously [33–35]. Mutant mice ([Tg/0; flox/flox]) together with their control littermates ([0/0; flox/flox] or [0/0; +/flox]) were housed in Plexiglas home cages (31.5 × 17 × 14 cm), each containing aspen bedding, nesting material, a wooden block for environmental enrichment, and 2–4 sibling animals. The access to food (standard rodent chow) and water was unlimited throughout the experiment. When mutant mice developed a pronounced motor phenotype, a small bowl of moist chow was added to provide easier access to food. The experiments were planned and executed in accordance with the ARRIVE guidelines [36] and European and Polish laws concerning the use and welfare of laboratory animals (Directive 2010/63/UE, European Convention for the Protection of Vertebrate Animals Used for Experimental and other Scientific Purposes ETS No. 123, and Polish Law Dz.U. 2015 poz. 266). All procedures were approved by the II Local Ethical Committee in Krakow (permits no. 197/2021 and 92/2022).

The mice were weighed before the first tamoxifen injection and then once weekly from 7 to 14 weeks after tamoxifen treatment. During the L-DOPA treatment phase (14–15 weeks after tamoxifen treatment), body weight was measured daily and used to calculate the L-DOPA dose. The data collected during the L-DOPA treatment phase were averaged in one week bins for statistical purposes. The general experimental schedule is summarized in Figure 1.

**Figure 1.**
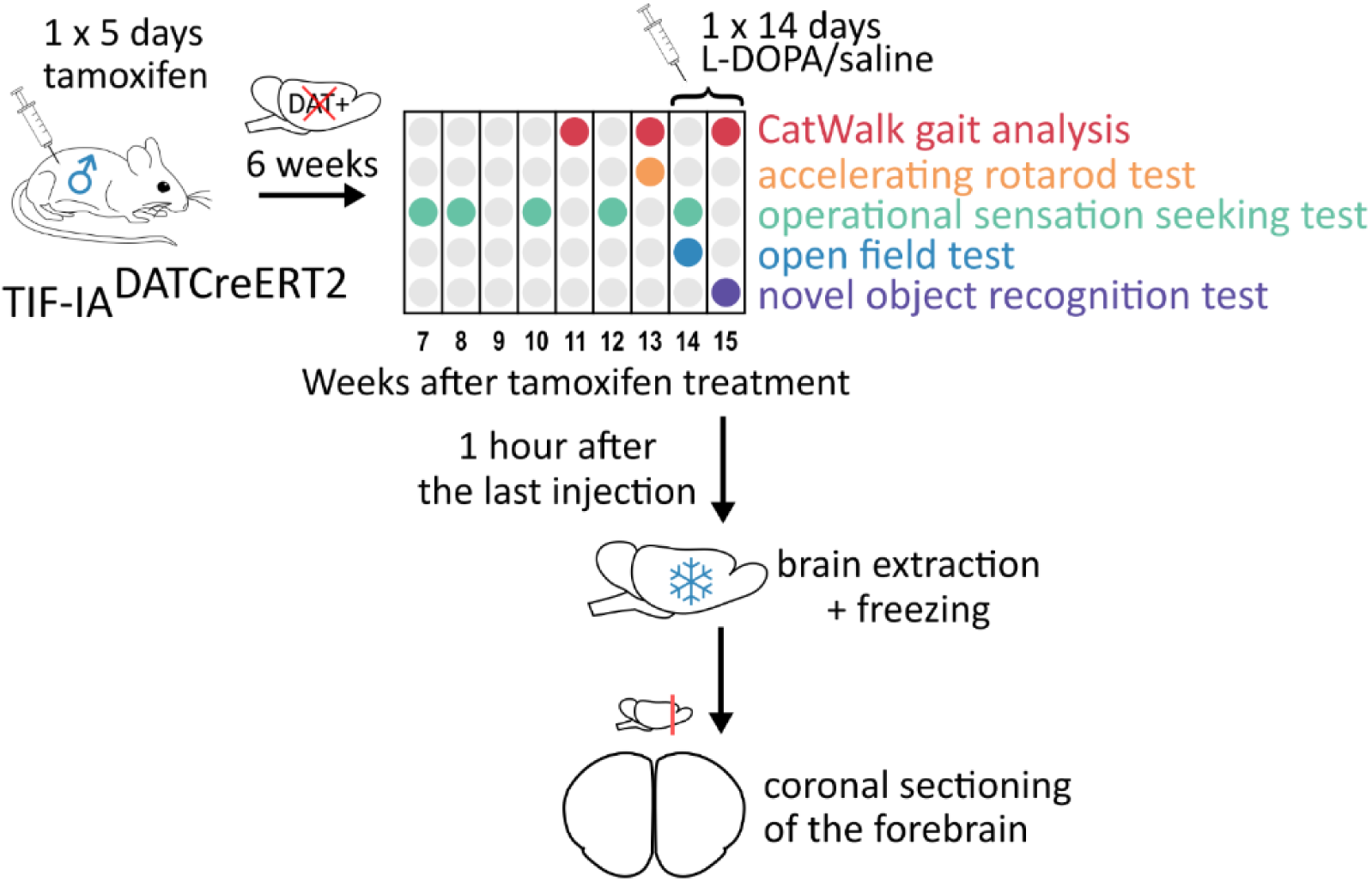
Experimental schedule. The diagram shows all major steps of the experiments performed: induction of the mutation with tamoxifen, behavioral phenotyping schedule, and brain preparation for further analyses.

### Drug treatment

Mutation was induced by i.p. tamoxifen administration (Sigma, Germany) daily for 5 consecutive days (2 mg of tamoxifen dissolved in 100 μl of sunflower oil) as described previously [35,37]. Starting from the 7th week after tamoxifen treatment until the end of the 15th week after the mutation was induced, the animals underwent behavioral testing of their motor abilities, cognitive function, and anxiety-like behaviors. During the 14th and 15th weeks after mutation induction, the mice were injected i.p. with benserazide hydrochloride (5 mg/kg, dissolved in saline) and 30 minutes later with L-DOPA (18 mg/kg in 10 μl, dissolved in saline by sonication; Sigma‒Aldrich, Germany). The control group consisted of mutant mice that were administered saline (Polpharma, Poland) in the volumes and intervals corresponding to the benserazide/L-DOPA treatments.

### Gait analysis

Gait features of mutant and control mice were assessed using the CatWalk XT system (Noldus, the Netherlands) 11, 13 and 15 weeks after tamoxifen treatment, as described previously [34,35]. Briefly, the mice were allowed to walk freely in a dark tunnel (130 × 20 × 0.5 cm), with green lighting below the glass floor. CatWalk XT v10.6 software was used to extract gait features. A total of 2–6 recordings of each mouse per test session were obtained. The selected measurements were averaged for both the front paws and hind paws as described previously [35].

### Accelerating rotarod test

Thirteen weeks after tamoxifen treatment, the mutant and control mice were tested using a Rotarod 47600 v04 apparatus (UGO BASILE, Italy), as described previously [34]. First, the mice underwent a training session, in which each mouse was placed on the apparatus for 5 minutes; the speed of the rotarod was constant at 6 rpm. The following day, the mice were placed on the rotarod at a constant speed of 6 rpm. After 20 s, the speed started to accelerate from 6 to 40 rpm over the course of 5 min, and the time to fall from the rod was recorded.

### Operant sensation-seeking test

This test was used to evaluate sensation seeking, memory, the ability to learn operant conditioning, and executive function in mice [38,39]. The procedure was performed as described previously and included training (daily for 5 consecutive days in the 7th week after tamoxifen treatment) and testing phases (3 times a week at 8, 10, 12, and 14 weeks after tamoxifen treatment) [35]. Briefly, the mice were placed individually in the soundproof conditioning chamber ENV-307 W (Med Associates, USA) for 60 minutes. The chambers were outfitted with two photocell-equipped holes placed on the wall opposite the cage light. When a mouse placed its snout inside the ‘active’ hole, the cage light was switched off, the diodes above the holes started blinking, and a monotone sound of 2.9 kHz was generated by an ENV-323AW Sonalert module. The blinking frequency was randomly selected from 0.625, 1.25, 2.5, or 5 Hz, and the signal was played for 2, 4, 6, or 8 s. The second hole was designated ‘inactive’, with no effects after the mouse’s action. The initial assignment of the active operant was randomized and then remained fixed throughout the experiment. The number of active and inactive nose pokes was recorded by MED-PC IV v4.2 software (Med Associates), and the results from individual sessions were averaged for each week.

### Novel object recognition test

A novel object recognition test was performed in the 15th week after tamoxifen treatment to examine short-term memory and recognition memory [40]. The procedure was performed according to a modified protocol described by Leger and colleagues [41] across 3 consecutive days (101, 102 and 103 days after the last tamoxifen dose) 60 minutes after the second L-DOPA (or saline) injection. The experimental room was dimly lit, with a radio playing in the background. The experimenter was not present in the room during the procedure. The animals were brought to the room ∼15 minutes before the experiment started. On the first day of the test, the mice were habituated to the test cage (37 × 21 × 18.5 cm, transparent, no bedding) for 5 minutes. On the second day, two identical objects were introduced into the cage: either two cell culture bottles (8 × 3.8 × 15.5 cm, transparent, partially filled with gravel) or two bottle-like shapes built from colored plastic blocks (6.2 × 3 × 15.5 cm; MegaBloks, Canada). The training session was 10 minutes. On the third day of the procedure, the mice were placed in the test cage for 10 minutes, with one familiar object replaced with a novel one. The placement of the novel object was counterbalanced. Animal behavior was recorded with an acA1300–60 gm (Basler, Germany) camera placed 1 m above the cage floor using EthoVision v11.5 software (Noldus). Recordings were analyzed with BORIS v7.13.6 software [42] to calculate the time spent exploring familiar and novel objects. Object exploration was defined as an event during which the mouse’s nose was directed at the object in the direct vicinity of it, and its behavior pointed clearly to the examination of the object and not the exploration of the cage. The recognition index was calculated as the fraction of time spent examining the novel object in relation to the total time spent examining both objects on the third day of the procedure. Animals whose total object exploratory time was shorter than 5 s were excluded from the analysis (3 animals in total).

### Open field test

An open field test was performed 14 weeks after tamoxifen treatment (1 hour after the second injection) to examine locomotor activity and anxiety-like behaviors [43,44]. The mice were placed in the center of an open arena (53 × 32 × 19 cm, opaque) lit with 10–12 lux light for 10 minutes. The animals were brought to the room ∼15 minutes before the experiment started. The room was dimly lit, with a radio playing in the background. The experimenter was not present in the room during the procedure. Mouse movement was recorded by an acA1300–60 gm camera placed 1 m above the cage floor using EthoVision v11.5 software. The recordings were analyzed with EthoVision v15.0 software.

### Spatial transcriptomic analysis of gene expression

The mice were euthanized by cervical dislocation 15 weeks after mutation induction and one hour after the last dose of L-DOPA (or saline) was injected. The brains were extracted, embedded in OCT medium (CellPath, United Kingdom), placed in 2-methylbutane, and flash-frozen in a liquid nitrogen bath. The brains were stored at -80 °C and sectioned using a CM 3050 S cryostat (Leica, Germany) with the following settings: a chamber temperature of -20 °C, an object temperature of -13 °C, and a section thickness of 10 μm. Sections were obtained from two regions: (1) the rostral part of the forebrain for spatial transcriptomic analyses (bregma 1.18 to 1.98 mm, as described previously [45]) and (2) the middle part of the forebrain to perform immunofluorescence staining for TH in the dorsal and ventral striatum (bregma 0.02 to 1.1 mm). The tissue blocks and microscopy slides were kept on the cryostat for at least 30 minutes before sectioning to avoid damage to the samples due to temperature differences. Sections intended for gene expression analyses were placed on Visium slides on one of 4 capture areas, each containing a matrix of spots with millions of oligonucleotide primers.

The spatial transcriptomic analysis was performed using Visium (10x Genomics, USA) in accordance with Visium Spatial Gene Expression User Guide Revision E. A total of 12 samples were analyzed with this method, using 3 Visium slides. Forebrain sections were first stained with hematoxylin and eosin and imaged separately in bright field using a Leica DMi8 system (Leica, Germany) and Leica Application Suite X v3.5.2.18963, together with an HC PL FLUOTAR 10x NA: 0.30 (DRY) objective and DFC 7000 camera. Images were processed with Loupe Browser v6.0 software (10x Genomics) to align the tissue position with the matrix of spots on Visium slides.

After imaging, the slide was fitted in a cassette, placed in a T-100 thermal cycler (Bio-Rad, USA), and permeabilized for 8 minutes. Next, the mRNA was reverse transcribed, and second-strand synthesis was performed according to the manufacturers’ instructions. cDNA was quantified using KAPA SYBR FAST qPCR Master Mix (Roche, Switzerland), amplified and then purified using SPRIselect paramagnetic beads (Beckman Coulter, USA). The quality and quantity of the cDNA were assessed with a high-sensitivity DNA kit (Agilent).

### Library construction and RNA sequencing

Illumina-compatible library construction was performed on the day after cDNA retrieval according to the manufacturers’ instructions. Briefly, 10 μl of each cDNA sample was fragmented (5 minutes at 32 °C), followed by end repair and A-tailing (30 minutes at 65 °C). SPRIselect was used for the double-sided size selection of samples, and cDNA was ligated with Illumina adaptors (15 minutes, 20 °C), purified, amplified, and labeled with i5 and i7 indices (Dual Index Plate TT Set A, 10x Genomics). The quality and quantity of the libraries were analyzed with a high-sensitivity DNA kit and the KAPA Library Quantification Kit Illumina Platforms (Roche). The libraries were sequenced by CeGaT GmbH (Germany) on a NovaSeq 6000 instrument (Illumina, USA) after equimolar pooling on SP or S1 flow cells. The following sequencing parameters were applied: 100 cycles and read lengths for Read 1 of 28 bp, for i7 of 10 bp, for i5 of 10 bp, and for Read 2 of 90 bp. The RNA sequencing parameters were selected according to the manufacturer’s recommendations to obtain approximately 125 million paired-end reads (250 million total reads) per sample. The capture areas varied between 36% and 67%, and the amount of the cDNA library per spot varied between 23.19 and 80.46 nM. The average fragment size in the cDNA libraries was between 407 and 438 bp. The total number of spots under the tissue in the 12 samples was 27,648. RNA-seq yielded ∼224 million average reads per library, corresponding to 46,161 paired-end reads per spot.

### Tyrosine hydroxylase (TH) immunostaining

Anti-TH staining was performed as described previously [35,46] with modifications. Briefly, 10 μm coronal sections from the same brain samples used to generate Visium libraries were used. Slides on which frozen sections were mounted were stored at -80 °C and transferred to dry ice after being frozen in 2-methylbutane in a liquid nitrogen bath. Each slide was fixed with 50 ml of chilled methanol (POCh) at -20 °C for 30 minutes, permeabilized in PBS with v/v 0.2% Triton X-100 (PBST) for 15 minutes at room temperature, and rinsed with PBS 3 times for 5 minutes. Next, they were incubated for 30 minutes with 5% v/v pig serum in PBST (Vector Laboratories), placed in Visium slide cassettes (10x Genomics) and incubated with an anti-TH antibody (AB1542, Sigma Aldrich) diluted 1:200 in blocking buffer overnight at 4 °C. Next, the slides were removed from the cassettes, rinsed with PBS twice for 5 minutes, then once for 15 minutes, and placed again in cleaned cassettes. The sections were incubated with secondary antibody (cat. no. A-11015, Invitrogen) at a ratio of 1:400 in blocking buffer for 1 hour at room temperature, rinsed three times for 5 minutes with PBS, and mounted with DAPI-containing medium (cat. no. AB104139, Abcam). Imaging was performed using a DMi8 system (Leica) in a fluorescent configuration coupled with an EL6000 lamp and a DAPI/FTC/TXR filter cube. Images were captured with a DFC 7000 camera (Leica) using Leica Application Suite X v3.5.2.18963 software and exported with gamma set to 0.5.

### Data analysis

Untrimmed, demultiplexed Reads 1 and 2 were quality-checked using fastQC v0.11.8 and then processed with the Space Ranger v1.3.1 (10xGenomics) software suite to align the reads to the mouse reference genome mm10-2020-A (GENCODE vM23/Ensembl 98), and the RNA-seq data were coupled with the spatial coordinates of each read. Further analysis was performed using a custom pipeline.

The resulting .bam files from all 12 samples were first merged using samtools [47] and processed using MACS3 (https://github.com/macs3-project/MACS) to identify and filter out peaks with too few reads. The reads were primarily annotated to gene or long terminal repeat (LTR) sequences at a distance of < 30 kbp from the corresponding peaks in both directions, together with the identification of the proper strand orientation. Next, we eliminated reads that were shown to be of uncertain genome alignment (107,651 total) from the analysis compared with data from a separate nanopore RNA-seq analysis of the striatum of two C57BL/6 mice [48]. The next step included filtering the peaks with the following criteria: amplitude > 600, number of reads assigned to a peak > 1200, number of reads assigned to amplitude ratio > 1.4, score > 350, summit.p.log > 35, and correct strand orientation (< 30 kbp in one direction from the gene/LTR sequence). Processed and annotated peaks were used to create a new reference transcriptome with the spaceranger mkref function, which was then used to perform a genome alignment of the RNA-seq data again with the spaceranger count command. The resulting transcript abundance data for each of the tissue spots across the 12 samples were then employed to perform a clustering analysis, visualize differences between the clusters, and perform a statistical analysis of the effects of L-DOPA.

The Seurat v4 package (https://satijalab.org/seurat/ [49]) was used to perform the unsupervised clustering analysis with the shared nearest neighbor method. Reads from 12 samples were used for clustering. The cluster resolution parameter was empirically set to 0.8 to ensure the optimal number of clusters that are compliant with different tissue domains in the mouse forebrain according to the Allen Mouse Brain Atlas [50]. Seurat’s UMAP plot and our custom-made functions were used to visualize the transcriptional profiles of each tissue spot. Cluster markers were identified with Seurat’s FindAllMarkers function with the min.pct parameter set to 0.4 and logfc.threshold set to 0.8. For each of the clusters, we estimated the false discovery rates (FDRs). One L-DOPA-treated mutant mouse with normal levels of the TH signal in the ventral and dorsal striatum was excluded from the statistical analysis but included in the unsupervised clustering analysis.

### Statistical analysis

The results of the behavioral analyses were assessed using analysis of variance (ANOVA). The results from the CatWalk test were assessed using a t test with the Benjamini‒Hochberg (BH) correction for the number of parameters assessed.

Spatial transcriptomic data were quantile-normalized. The effects of L-DOPA were determined for mutants only, i.e., 5 treated with saline and 4 treated with L-DOPA, with t tests performed for each cluster separately. The dataset included 15,955 transcripts corresponding to 10,545 genes with Ensembl IDs. The significance criterion was set as p < 0.01, log2 ratio ≥ 0.5, and mean normalized abundance ≥ 0.6 in at least one of the groups. All the statistical analyses were performed in R.

## RESULTS

### Phenotypes of the TIF-IA^DATCreERT2^ mice

The induction of the mutation caused the development of a phenotype resembling some PD symptoms. As reported previously [35], TIF-IA^DATCreERT2^ mice presented reduced weight gain, with the largest effect observed at 14 weeks after induction of the mutation (Figure 2a; F_1,22_ = 4.55, p = 0.0443 time: F_8,176_ = 46.77, p < 0.0001; time × genotype F_8,176_ = 10.46, p < 0.0001). The difference in weight persisted throughout the L-DOPA treatment period, with a mean weight of mutant mice of 27.8 ± 0.939 g compared with 31.9 ± 0.655 g in the controls (Figure 2a; two-way ANOVA with interaction: genotype: F_1,19_ = 13.823, p = 0.00146; treatment: F_1,19_ = 0.867, p = 0.36359; genotype × treatment: F_1,19_ = 0.704, p = 0.41195). At 13 weeks after the induction of the mutation, the TIF-IA^DATCreERT2^ mice presented a significant motor impairment in the rotarod test; compared with the control mice, the mutant mice remained on the accelerating rod for 37 s on average, whereas the control mice remained on the accelerating rod for 140 s (Figure 2b). These results are consistent with previous reports on rotarod performance after an 80% loss of dopaminergic neurons in the SNpc of TIF-IA^DATCreERT2^ mice 3 months after mutation induction[33–35].

**Figure 2.**
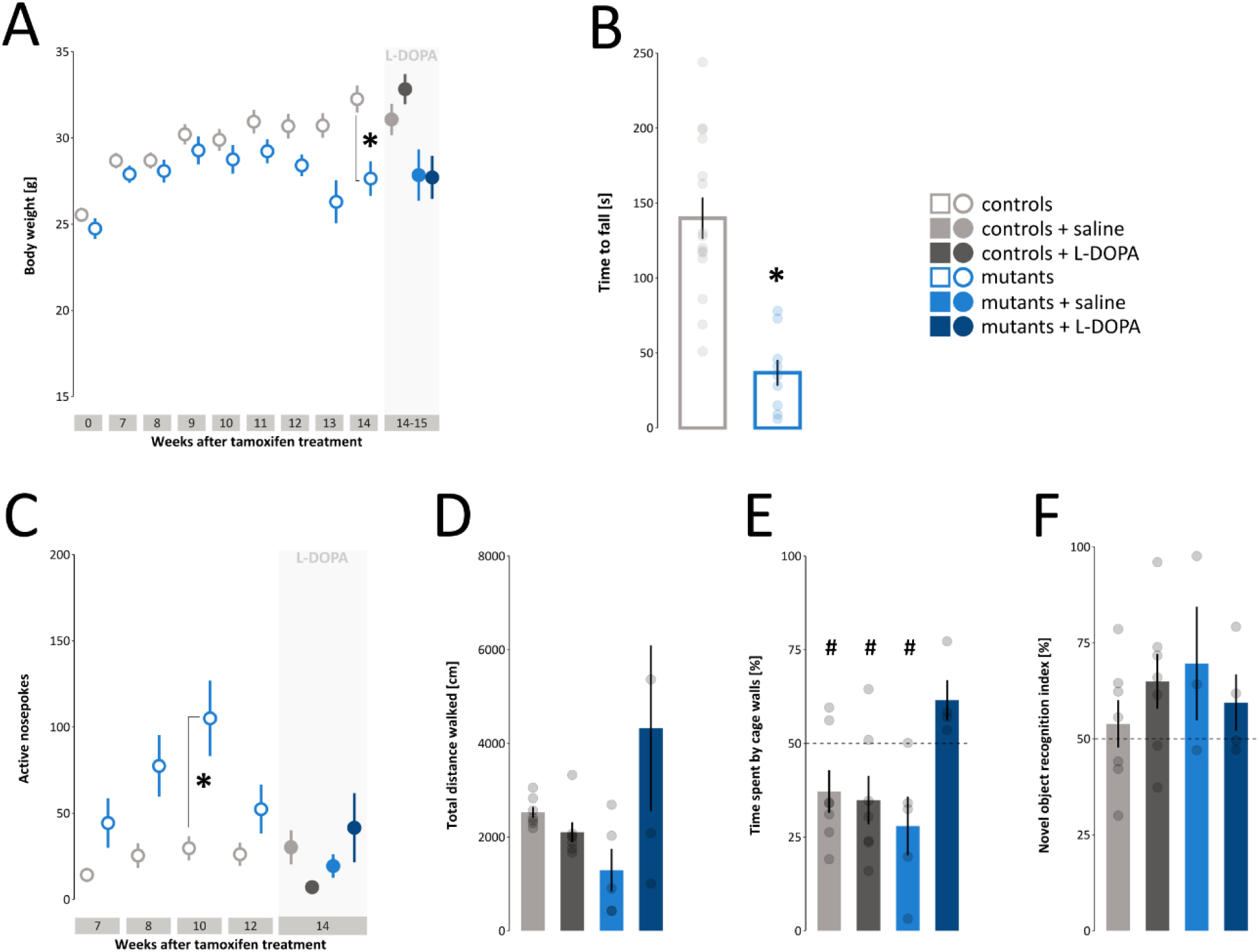
Phenotype of the progressive loss of dopaminergic neurons in adult male TIF-IA^DATCreERT2^ mice. (a) Reduced weight gain in TIF-IA^DATCreERT2^ mice. Each point represents the mean weight at a time point relative to induction of the mutation, with “0” being the measurement taken just prior to tamoxifen treatment. The area shaded in gray represents the period corresponding to L-DOPA treatment. (b) Mean time that the animals were able to remain on the accelerating rotarod during the test performed 13 weeks after the induction of the mutation. (c) Mean number of instrumental responses to the active operant in the operant sensation-seeking test. Tests were performed at specific time points after induction of the mutation, as shown below the graph. The area shaded in gray represents the period corresponding to L-DOPA treatment. (d) Mean distance traveled by the mice in the open field at 14 weeks after mutation induction and during L-DOPA treatment. (e) Mean time the mice spent in the central zone of the open field in the same experiment as described D. (f) Mean recognition index in the novel object recognition task performed at 14 weeks after mutation induction and during L-DOPA treatment. The control animals are shown in gray, the mutant mice are shown in blue, and the color shading corresponds to the treatment, as shown in the legend in the upper right. The error bars represent s.e.m.s. Individual measurements are shown in all the bar graphs.

The progressive loss of dopaminergic neurons caused changes in static and dynamic gait parameters. A significant effect of the mutation was observed, including a larger front paw support base (Table S1, genotype: F_1,21_ = 9.795, p = 0.00506; time: F_1,21_ = 3.423, p = 0.08; time × genotype: F_1,21_ = 0.023, p = 0.88) and the decreased coupling between the front and hind right paws (genotype: F_1,21_ = 6.122, p = 0.022; time: F_1,21_ = 0.005, p = 0.94; time × genotype: F_1,21_ = 1.45, p = 0.24). Altered gait parameters persisted during L-DOPA treatment, including couplings between the front and hind right paws (Table S1, genotype: F_1,19_ = 6,331, p = 0.021; treatment: F_1,19_ = 0.91, p = 0.352; genotype × treatment: F_1,19_ = 0.84, p = 0.371) and the mean front paw stride length (genotype: F_1,19_ = 6.078, p = 0.0234; treatment: F_1,19_ = 3.04, p = 0.1; genotype × treatment: F_1,19_ = 0.017, p = 0.9). Additionally, significant interaction effects between the treatment and genotype on the mean duration of the front paw step cycle (genotype: F_1,19_ = 0.73, p = 0.404; treatment: F_1,19_ = 1.42, p = 0.25; genotype × treatment: F1,19 = 10.46, p = 0.004), cadence (genotype: F_1,19_ = 0.15, p = 0.7; treatment: F_1,19_ = 0.42, p = 0.526; genotype × treatment: F_1,19_ = 9.43, p = 0.00629), and base of support for hind paws (genotype: F_1,19_ = 4.14, p = 0.056; treatment: F_1,19_ = 0.02, p = 0.89; genotype × treatment: F_1,19_ = 6.484, p = 0.0197) were observed. However, these effects were not corrected for the number of parameters tested. Despite the motor impairments, at 14 weeks after mutation induction, the mutant mice showed no reduction in locomotor activity. L-DOPA treatment caused an increase in motor activity in mutant mice in terms of both the distance traveled and movement speed (Figure 2c, distance: genotype: F_1,19_ = 0.250, p = 0.623; treatment: F_1,19_ = 2.071, p = 0.166; genotype × treatment: F_1,19_ = 6.989, p = 0.016; speed: genotype: F1,19 = 0.253, p = 0.621; treatment: F_1,19_ = 2.085, p = 0.1651; genotype × treatment: F_1,19_ = 7.012, p = 0.0159). Notably, L-DOPA treatment increased the proportion of time that mutant mice spent in the outer zone of the field, which indicated an increase in anxiety-like behaviors (Figure 2d, genotype: F_1,19_ = 1.038, p = 0.3210; treatment: F_1,19_ = 3.129, p = 0.0930; genotype × treatment: F_1,19_ = 7.113, p = 0.0152).

Furthermore, TIF-IA^DATCreERT2^ mice developed nonmotor behaviors consistent with our recently reported findings [35], including an increase in instrumental responses in the operant sensation-seeking test (Figure 2e; genotype: F1,22 = 11.18, p = 0.00294; time: F3,66 = 10.526, p < 0.00001; genotype × time: F3,66 = 6.703, p = 0.000512). L-DOPA treatment further increased the number of instrumental responses in mutant mice (Figure 2e; genotype: F1,19 = 1.137, p = 0.2996; treatment: F1,19 = 0.328, p = 0.5736; genotype × treatment: F1,19 = 5.214, p = 0.0341). Conversely, memory in the novel object recognition test was not affected by the mutation (Figure 2f, genotype: F1,17 = 0.265, p = 0.613; treatment: F1,17 = 0.257, p = 0.619; genotype × treatment: F1,17 = 1.545, p = 0.231). Taken together, the TIF-IA^DATCreERT2^ mice displayed specific behaviors similar to PD symptoms, including weight loss and a moderate motor impairment. Additionally, TIF-IA^DATCreERT2^ mice performed a greater number of instrumental responses in an operant task and had increased sensitivity to L-DOPA than controls did.

### Spatial transcriptomics

A spatial gene expression analysis was performed on coronal brain sections obtained from 5 mutant mice treated with saline, 5 mutant animals treated with L-DOPA, and 2 controls treated with saline that were euthanized 15 weeks after mutation induction and 1 h after the final drug injection. First, sections from selected brains were stained to determine the expression of tyrosine hydroxylase (TH) in the basal ganglia and validate the efficiency of the mutation. As shown in the representative images in Figure 3, a robust anti-TH signal was observed in the caudate putamen, nucleus accumbens and olfactory tubercle of a control animal (Figure 3a) but was greatly reduced or absent in sections derived from mutant mice (Figure 3b and c), with the remaining signal most visible in the ventromedial area. One mutant animal was excluded from further analyses because of the high control level of the anti-TH signal.

**Figure 3.**
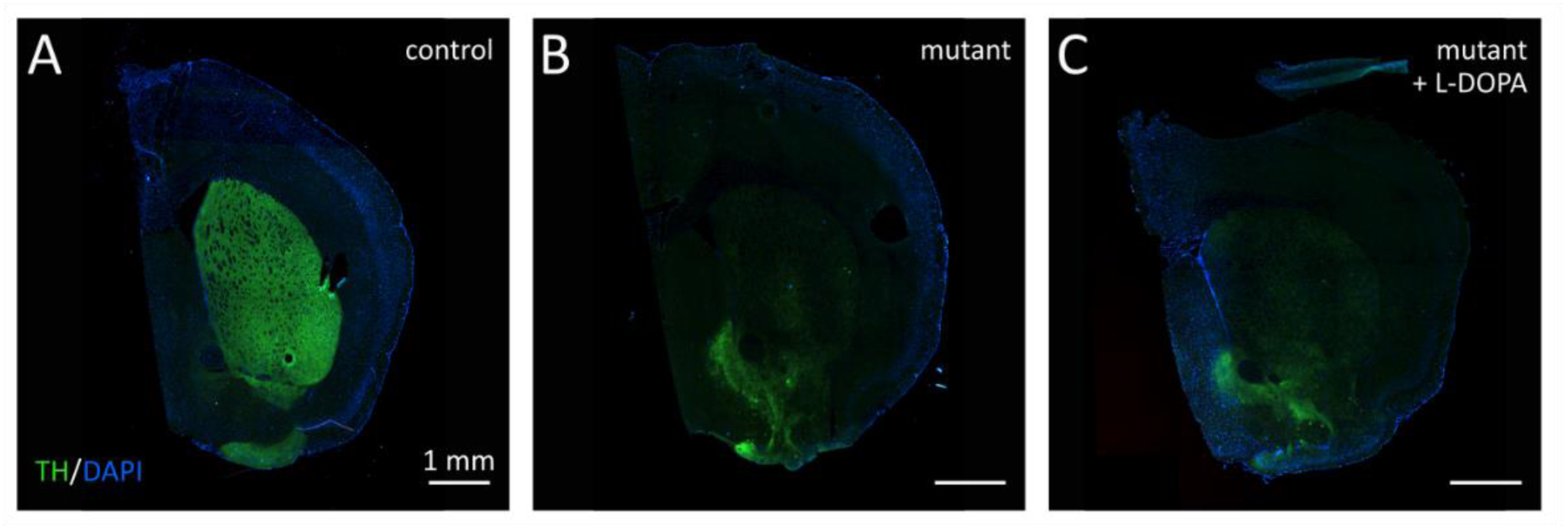
Loss of dopaminergic fibers in the striatum of TIF-IA^DATCreERT2^ mice. Representative micrographs showing tyrosine hydroxylase (TH) immunofluorescence staining (green) in coronal sections from (a) control, (b) mutant, and (c) L-DOPA-treated mutant mice. The cell nuclei were stained with DAPI (blue). Scale bars are shown in the lower left corners of the panels.

Next, cDNA libraries were generated from sections transferred onto Visium slides, sequenced, and aligned to the mouse reference genome mm10-2020-A. The initial analysis revealed multiple cases where reads corresponding to mRNA 3’ termini were mapped outside known transcripts, usually further downstream from annotated ends [48]. We developed a custom pipeline that additionally checks for the correct strand match between the transcript and reads to minimize the impact of mismatches. First, the spaceranger count function was used to generate .bam files, and 24,209 unique genes corresponding to 24,224 Ensembl IDs were identified. Next, the experiment-wide .bam files from all 12 samples were merged, and the SAMtools package was used to identify 173,758 read ‘peaks’. The MACS3 package was used to analyze the shapes of the peaks and exclude alignments to repeated sequences. We subsequently used data from previously reported long-read sequencing of mRNAs derived from a coronal mouse brain section corresponding to the same area as that assessed in spatial analyses [48] to validate the matches. A total of 157,803 reads were excluded based on the custom reference, and the great majority represented a single read mapped to a transcript or reads not matching long-read sequences. The procedure yielded 15,955 peaks corresponding to genes with Ensembl IDs, and this set was used for transcript counting and all further analyses.

An unsupervised clustering analysis was performed on all the spots using the Seurat software package v4 and the shared nearest neighbor method, with the resolution parameter set to 0.8, which resulted in 27 clusters (Figure 4a). The clustering results accurately recapitulated the layers in the dorsal (#1, #3, #4, #5, #6, #7, and #18) and piriform (#17 and #21) cortices, closely matching a schematic representation from the Allen Reference Atlas–Mouse Brain [50] (Figure 4b). Dorsal cortical layers 6a and 6b were separated into two substructures (#7 and #22, respectively), clusters #5 and #1 matched layers 5a and 5b of the neocortex, and cluster #6 corresponded to neocortical layer 5. The claustrum and endopiriform nucleus were grouped together as cluster #14. Remarkably, the analysis clearly separated functional areas of the basal ganglia: the dorsal striatum (#0), the nucleus accumbens (#2), and the olfactory tubercle (#11). Scattered spots probably corresponding to fiber tracts in the dorsal striatum were grouped separately (#25), and ventrolateral parts of the accumbens and striatum were separated into their own substructures (#24 and #11, respectively). Areas predominantly composed of white matter (#10) and meninges (#8) were grouped separately. As shown in the UMAP projection of the clustering results (Figure 4c), the expected similarities in the transcript profiles of spots from areas rich in medium spiny neurons (left side, #0, #2, and #11), lower cortical layers (lower right, #1 and #7) and upper cortical layers (upper right, #3 and #4) were observed. Furthermore, the anatomical overlap between the clustering results and brain structures was reflected in the presence of the expected transcript markers in specific brain areas, including *Tac1* and *Penk* in the dorsal striatum (Figure 5a, #0, #2, #11, #24, and #25, [51,52]), *Lamp5* and *Rasgrf* in cortical layers 2/3 (Figure 5b, #3, [53]), or *Lxn* and *Gng2* in cortical layer 6 (Figure 5c, #14, [54]). Notably, the expression of individual marker genes is not exclusive to the corresponding area, and in particular, in the case of the cortical markers, they are ubiquitously present in the cortex, albeit at lower abundances. Rather, brain substructures were defined by extended sets of markers, and these patterns provided a highly accurate reconstruction of the brain anatomy.

**Figure 4.**
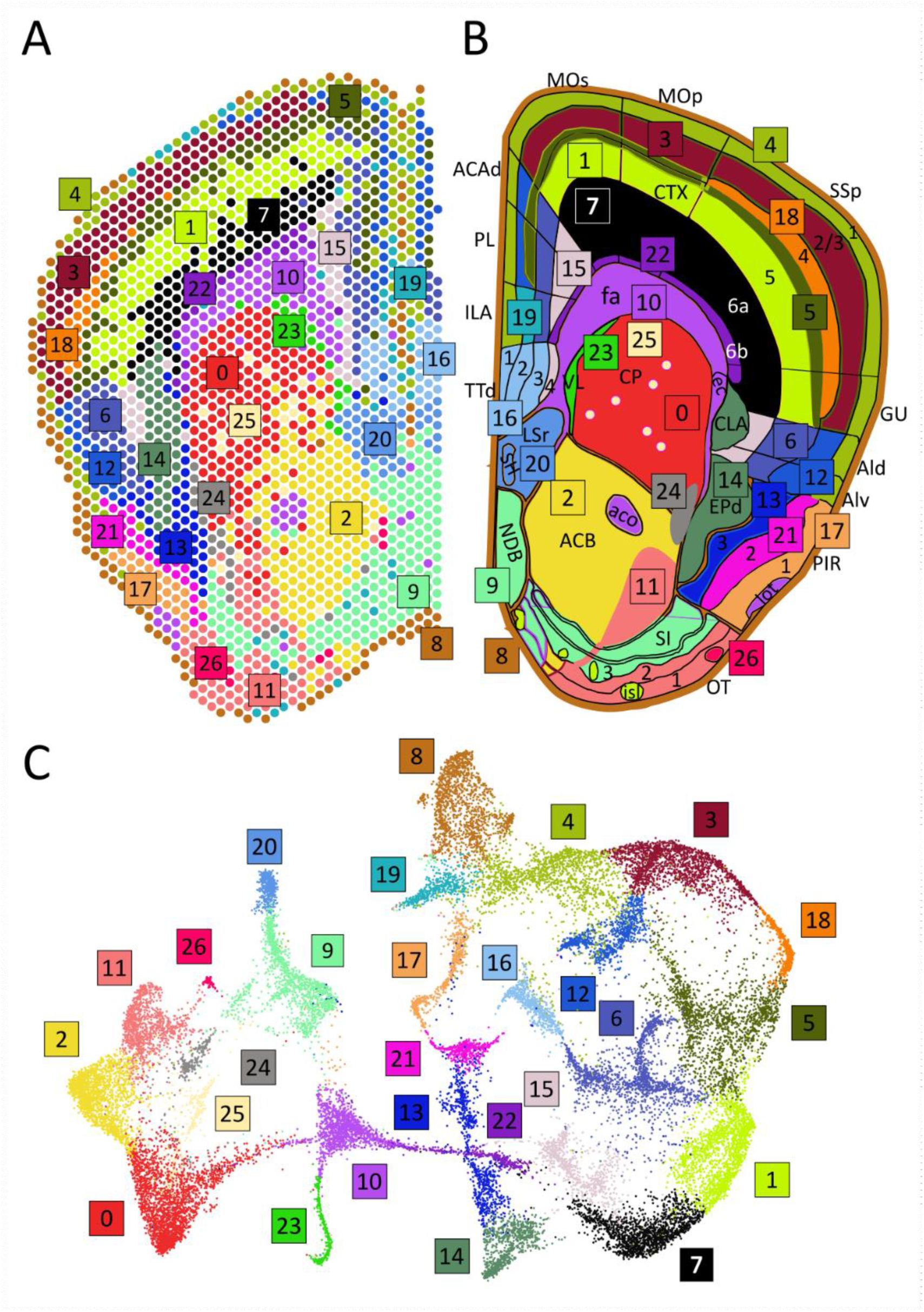
Clustering analysis of the spatial transcriptomic data. (a) Distribution of clusters projected on a coronal section of a brain hemisphere. Each sequenced transcript was mapped to one spot. The colors are used to represent 27 clusters, with the corresponding numbers (starting from #0) shown in colored boxes. (b) A schematic representation of brain structures adapted from the Allen Mouse Brain Atlas; the colors mark areas corresponding to the clusters shown in (a). (c) UMAP projection of the clustering of all individual profiles (“spots”) from all sections. The numbers and colors correspond to Panels (a) and (b). Abbreviations: ACAd, anterior cingulate cortex, dorsal part; ACB, nucleus accumbens; aco, anterior commissure; Ald, agranular insular area, dorsal part; Alv, agranular insular area, ventral part; CLA, claustrum; CP, caudoputamen; CTX, cortex; EPd, endopiriform nucleus, dorsal part; fa, corpus callosum, anterior forceps; GU, gustatory areas; ILA, infralimbic area; isl, islands of Calleja; lot, lateral olfactory tract; LSr, lateral septal nucleus, rostral part; MOp, primary motor area; MOs, secondary motor area; NDB, diagonal band nucleus; OT, olfactory tubercle; PIR, piriform area; PL, prelimbic area; SSp, primary somatosensory area; SH, septohippocampal nucleus; SI, substantia innominata; TTd, taenia tecta, dorsal part; VL, lateral ventricle.

**Figure 5.**
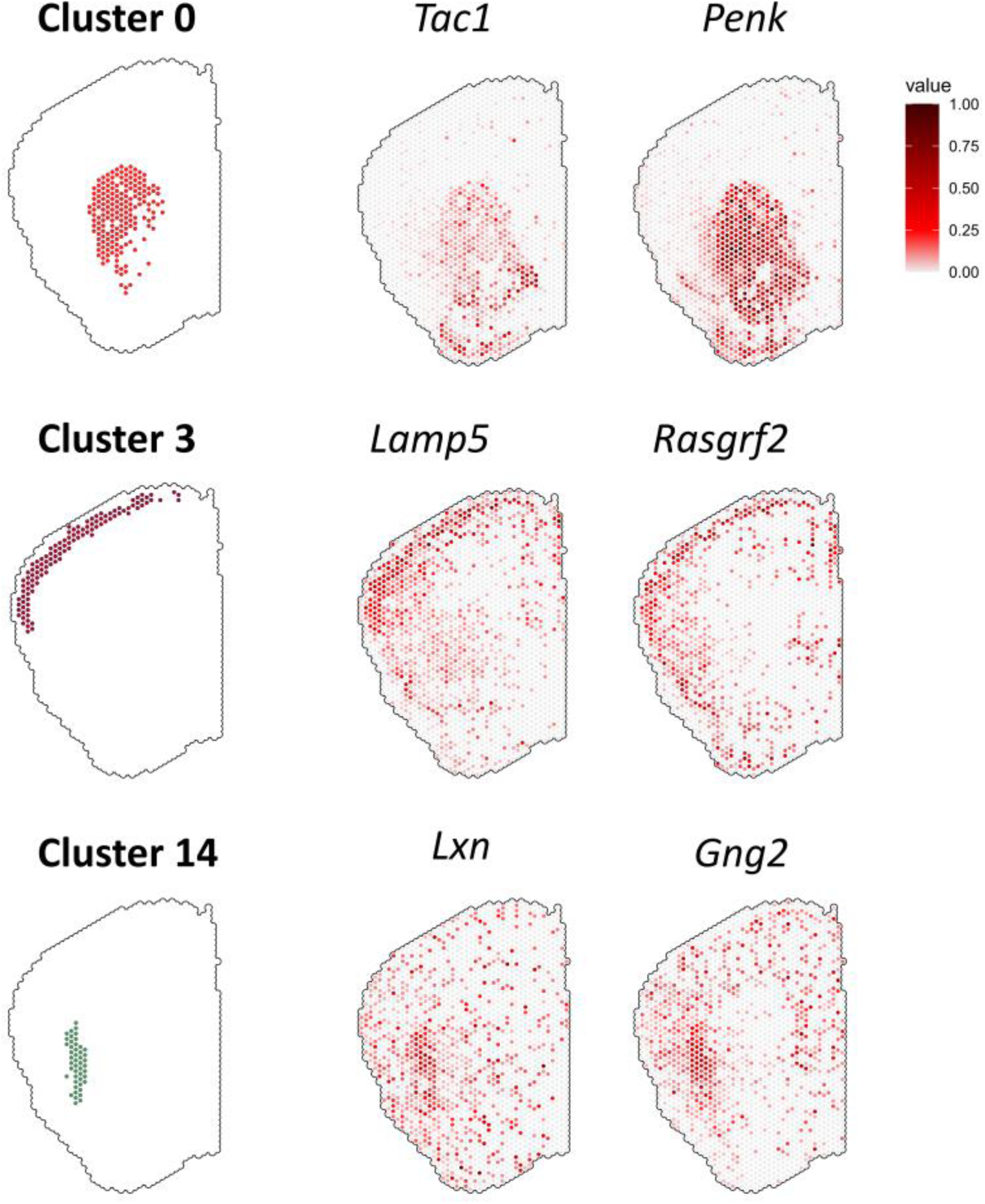
Representative profiles of cluster-defining transcripts. Examples of 3 clusters (#0, #3 and #14) and their distributions in a representative section. For each cluster, two examples of defining transcripts are given: *Tac1* and *Penk* for #0, *Lamp5* and *Rasgrf2* for #3, and *Lxn* and *Gng2* for #14. In the case of the transcripts, each dot represents the relative abundance according to the legend shown in the upper right.

### L-DOPA-regulated gene expression

The effects of L-DOPA on transcription in TIF-IA^DATCreERT2^ mice were assessed within each structure, and the significance criterion was set as p < 0.01, log2 ratio ≥ 0.5, and mean normalized abundance ≥ 0.6. Significant drug-induced changes in gene expression were observed for 181 transcripts corresponding to 121 genes (88 whose expression increased and 33 whose expression decreased) in 23 clusters, as summarized in Figure 6. The largest group of L-DOPA-induced genes was activity-dependent genes, including *Arc*, *Bhlhe40*, *Dusp1*, *Dusp6*, *Egr1*, *Fos*, *Fosl2*, *Homer1*, *Junb*, *Nr4a1* and *Nr4a2*. These changes were observed in the dorsal striatum (#0) as well as layers 2/3 and 5 of the dorsal cortex (#3 and #1, respectively) and were generally widely expressed in the case of transcripts such as *Egr1* (13 structures with significant increases in transcript abundance), *Nr4a1* (in 9 structures) or *Junb* (in 5 structures, Figure 7). Notably, while trends toward an increase in the expression of several of these transcripts were observed in the nucleus accumbens (#2), none of them reached significance. The differences and similarities between gene expression patterns between layers 2/3 of the cortex, striatum and nucleus accumbens were clearly visible when normalized transcript levels were compared (Figure 8). For example, no appreciable induction of the expression of *Fos*, *Egr1* or *Junb* was detected in the nucleus accumbens (#2), and the expression of transcripts such as *Bhlhe40* was selectively induced in dorsal cortical layers 2/3 (#3), whereas *Nptx2* (Figure 7b) or *Adam23* was upregulated in the striatum (#0). Finally, the expression of transcripts such as *Jup* (junction plakoglobin) appeared to be consistently induced in all three areas, even though these effects did not reach the significance criterion in the nucleus accumbens. Thus, the spatial pattern of L-DOPA-induced changes in gene expression was heterogeneous, with specific transcripts showing ubiquitous or highly structure-specific changes.

**Figure 6.**
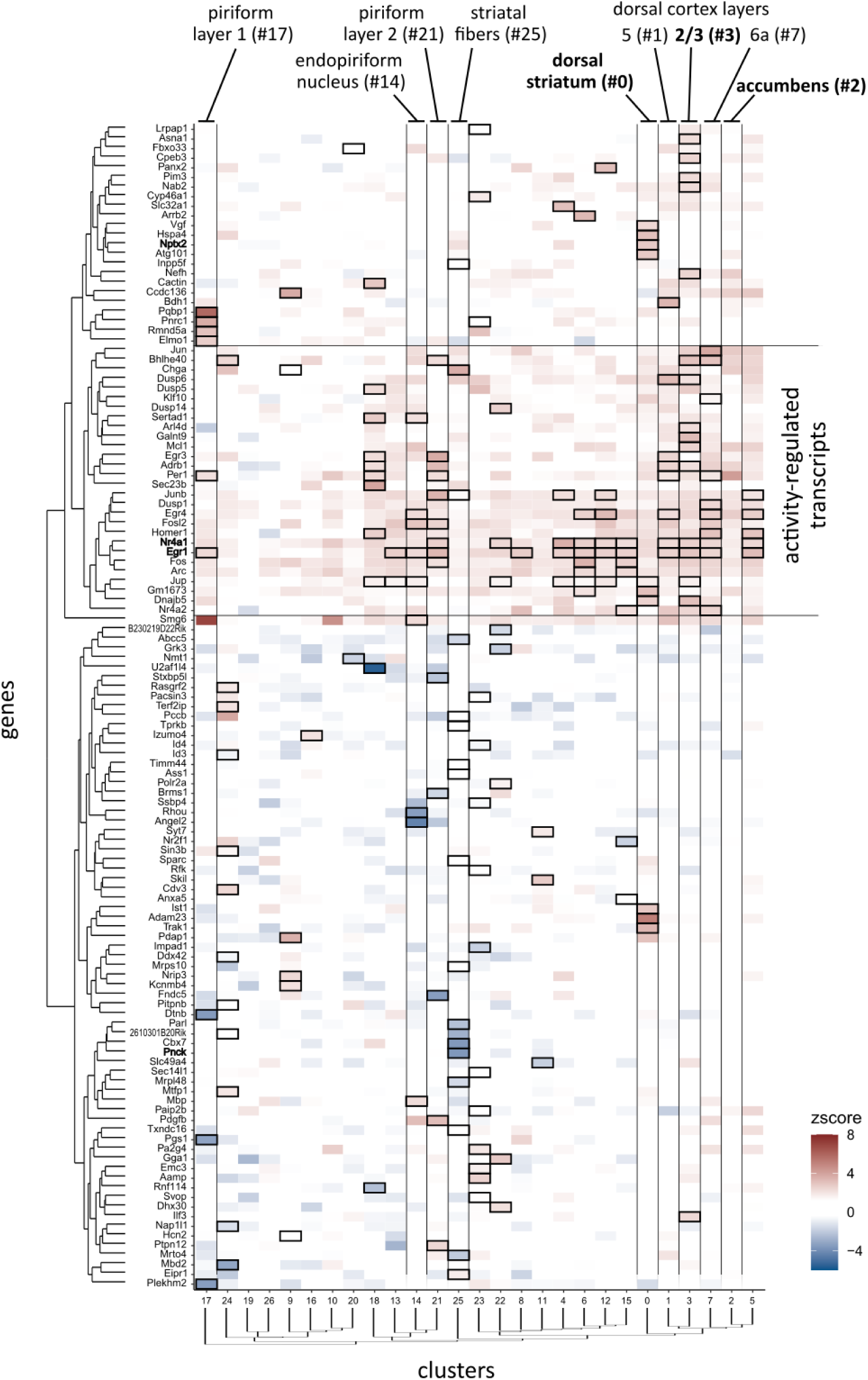
L-DOPA-induced gene expression in TIF-IA^DATCreERT2^ mice. The heatmap summarizes all significant changes in gene expression caused by L-DOPA treatment in all clusters. Each column represents the z score of the difference between mutant mice treated with L-DOPA and saline-treated mutant mice; the color legend is shown in the lower right corner, and cluster numbers are shown at the bottom. Each row represents one gene indicated on the left. Boxes indicate a significant effect of L-DOPA on transcript abundance in a specific cluster. The genes and clusters were ordered by clustering, as shown in the dendrograms on the left and below the heatmap. Two horizontal lines mark a group of genes that were designated activity-regulated transcripts. Vertical lines were added to mark the clusters indicated above the heatmap. The cluster names shown in bold correspond to the 3 clusters shown in Figure 8, and the gene names shown in bold correspond to the transcripts shown in Figure 7.

**Figure 7.**
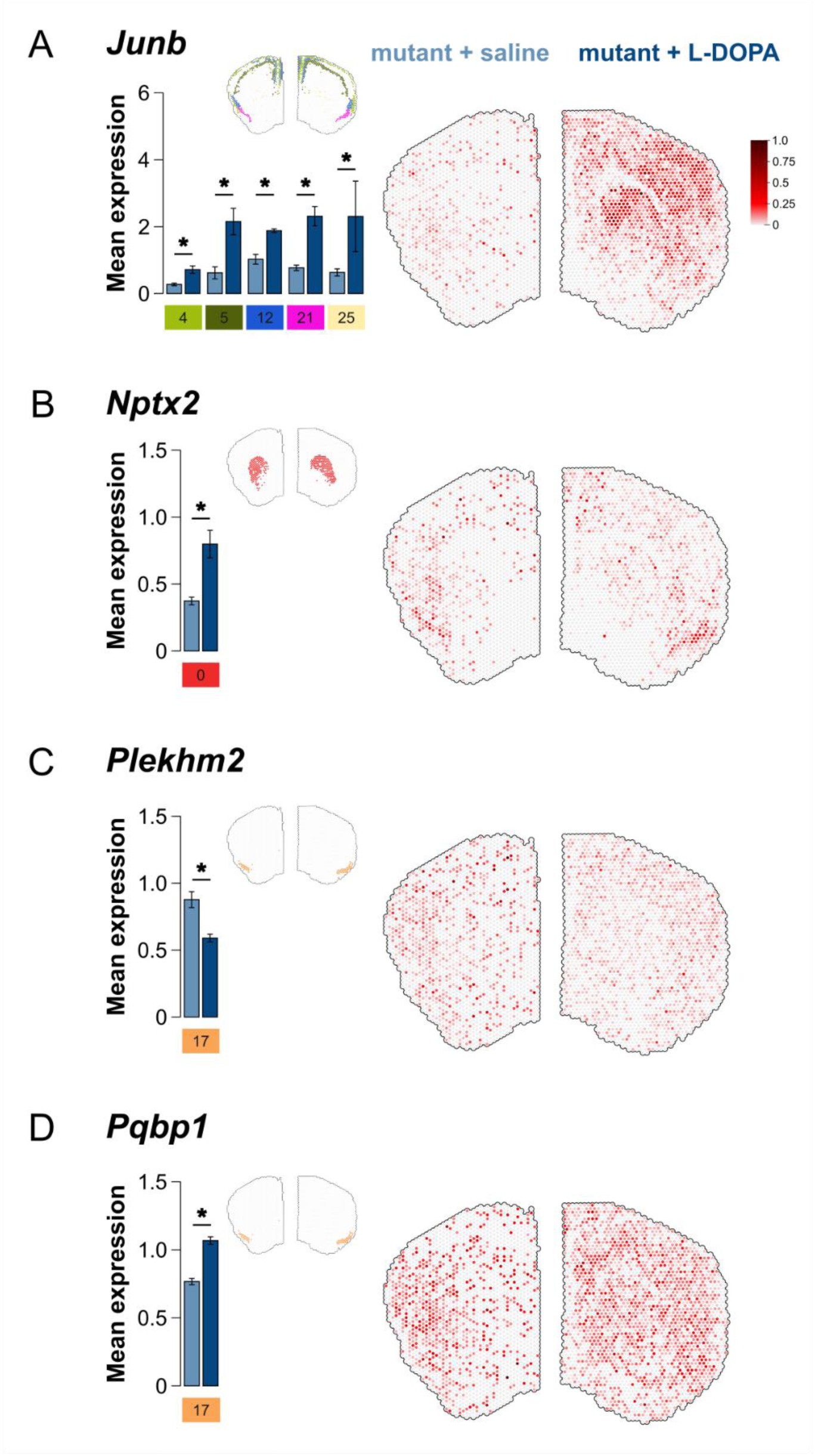
Representative examples of L-DOPA-regulated transcripts: (a) *Junb*, (b) *Nptx2*, (c) *Pqbp1*, and (d) *Plekhm2*. Each panel shows a representative example of the transcript distribution in sections corresponding to a control mouse and L-DOPA-treated TIF-IA^DATCreERT2^ mouse (right panel), a bar graph showing significant changes in expression in selected clusters (left panel), and the distribution of the clusters on the representative sections (above the bar graph on the right). The bars represent the mean normalized values ± s.e.m.. The legend in the upper right shows the color intensity corresponding to the normalized expression level.

**Figure 8.**
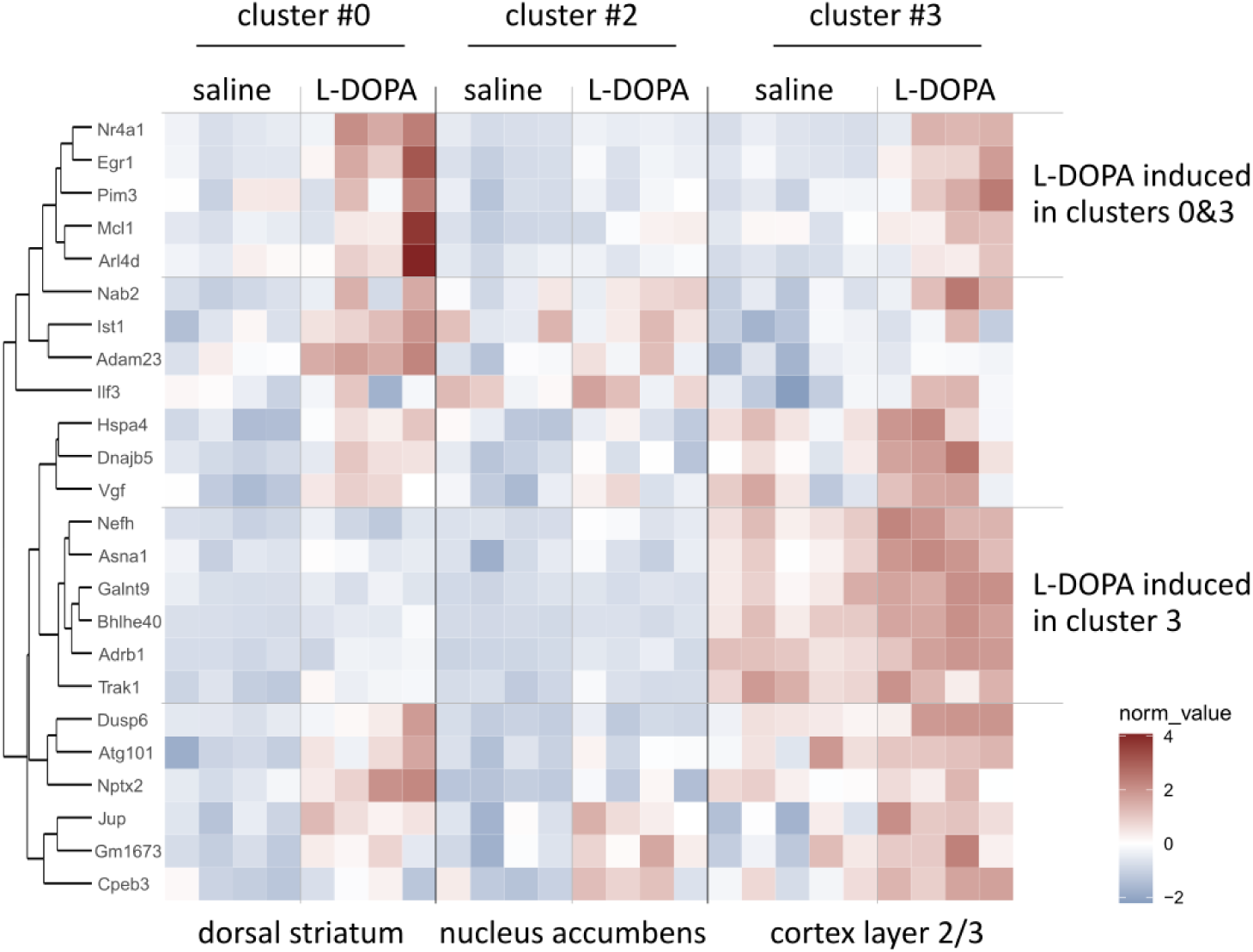
Differences in L-DOPA-induced gene expression in the striatum, nucleus accumbens and cortical layers 2/3. The heatmap shows normalized levels of expression, and the color scale is shown at the bottom right. Each row represents a transcript, as indicated on the left. Transcripts were clustered as shown in the dendrogram on the left. Each column represents a single animal/section, grouped according to cluster and treatment, as shown above the heatmap. Vertical lines were added to mark groups of transcripts induced in clusters #0 and #3 or selectively in cluster #3.

We found that L-DOPA treatment also caused significant decreases in the abundance of transcripts, including *Plkhm2* (Figure 7c), *Pnck, Dtnb* and *Angel2*. However, in none of the cases were these changes significant in more than one cluster; with few exceptions, they were detected in three areas corresponding to the piriform cortex (#17 and #21), the endopiriform nucleus and claustrum (#14), and a diffuse group of spots that resembled the white matter fibers crossing the dorsal striatum (#25, the eponymous “stripes”). Notably, no significant decreases in transcript abundance were detected in the striatum (#0), nucleus accumbens (#2), or dorsal cortex (#1, #3, #5, and #7). Conversely, the expression of a group of transcripts that was specifically induced selectively in the upper layer of the piriform cortex (#17) included *Pqbp1* (Figure 7d), *Pnrc1*, *Rmnd5a*, and *Elmo1*. Moreover, in the piriform cortex and adjacent areas, increases in activity-dependent gene expression (e.g., *Junb*) were observed, which further highlights the ubiquity of this pattern. Finally, the difference in the expression pattern between cluster #25 and the surrounding striatum (#0) is also striking, as only spatial profiling is capable of detecting such effects.

The ontological analysis of differentially regulated genes showed a significant enrichment of GO terms linked to DNA-templated transcription (24 genes, GO:0006355), miRNA transcription (4 genes, GO:1902893), chemotaxis (GO:0060326), peptidyl-tyrosine dephosphorylation (GO:0035335), positive regulation of tau-protein kinase activity (GO:1902949) and 18 additional terms listed in Table S2. Additionally, based on the DisGeNET 2023 database of disease-related genetic variants [55], a significant enrichment in genes linked to 56 disease-relevant terms, including PD, depressive disorders, visual seizures, cocaine dependence, schizophrenia, neoplasms of various organs, and cardiovascular events (see the full list in Table S3). These results are consistent with the induction of activity-regulated transcription, most of which involve immediate-early genes that act as transcription factors (e.g., *Egr1*, *Junb* or *Fos*). Among the 16 differentially expressed genes linked to PD was *Nptx2*, which was selectively induced in the striatum. Taken together, the results of the functional analysis of the differentially expressed genes mostly reflect the involvement of the majority of the activity-regulated transcripts in transcription.

## DISCUSSION

The spatial analysis of L-DOPA-induced gene expression reveals the complexity of its actions in detail. We show that L-DOPA treatment has widespread effects extending beyond the primary areas involved in dopamine-dependent control of movement, with both ubiquitous and region-specific effects on transcription. A major difference between the findings of this study and those of previous studies on the effects of L-DOPA on rodents with lesions targeting dopaminergic neurons is the choice of model. To the best of our knowledge, all previous reports on L-DOPA-induced gene expression profiles are based on extensive and rapid 6-OHDA- or MPTP-induced dopaminergic neuron degeneration. The TIF- IA^DATCreERT2^ mouse model presented here is characterized by a slow, progressive loss of dopaminergic cells that starts in adulthood, which is caused by tamoxifen-induced cell type-specific loss of TIF-IA, an essential RNA transcription factor [33–35]. Notably, at the time point selected for gene expression, 14 weeks after the mutation was induced, some dopaminergic fibers were still present in the striatum, particularly in the ventral region (as shown in Figure 3 and previously reported [34,35]). Accordingly, the effects of L-DOPA on the expression of activity-regulated genes in the nucleus accumbens (#2 and #11) were limited and did not reach significance, although they exhibited a trend similar to that observed in the dorsal striatum (#0). Moreover, while the TIF-IA^DATCreERT2^ mice showed significant movement impairments at 14 weeks, the animals did not yet suffer from catalepsy. This stage of the progressive loss of dopaminergic neurons was selected to model the typical condition in humans, where pharmacotherapy is usually applied after early motor symptoms are observed. Notably, the effects of PD are not limited to the dopamine system; in particular, nonmotor symptoms may be influenced by the degradation of other types of neurons (e.g., cholinergic neurons in the case of cognitive symptoms, [1]). Thus, the analysis presented here shows the effects of two weeks of L-DOPA treatment on mice with an advanced but incomplete loss of dopaminergic neurons.

Consistent with previous studies, we found that L-DOPA induced the expression of activity-regulated genes in the dorsal striatum, including *Fos*, *Fosb*, *Junb*, *Arc* and *Egr1,* e.g., [26,28–31]. In some cases (e.g., *Fos*, *Egr1* or *FosB*), L-DOPA-induced expression was also previously observed via in situ hybridization or immunohistochemistry in the dorsal cortex, e.g., [18,20]. To a limited extent, the induction of activity-regulated expression was also observed in our previous study on the effects of L-DOPA on the prefrontal cortex of 6-OHDA-lesioned rats, where we observed the induction of *Per1* expression*, among other genes* [35]. However, these effects appeared minor compared with the changes observed here. This finding is consistent with weaker or no effects of L-DOPA on the medial cortex (#16 and #19) and may also be related to methodological differences: a different model organism (a rat), a different method of dopaminergic neuron lesion (unilateral 6-OHDA lesion), and an L-DOPA dosage of 12.5 mg/kg. Finally, the induction of activity-regulated transcript expression was also observed in the piriform cortex, particularly the cluster corresponding to layer 2 (#21), which was also observed in one of the early reports on L-DOPA-induced transcription [19]. The broad induction of activity-regulated transcripts is not specific to L-DOPA, and a similar pattern is observed when naïve animals are treated with psychostimulants or drugs of abuse [56,57]. Moreover, it appears to be a general response to drug treatment [58–60] and was also observed in the case of behavioral stimuli, including exposure to stress, a novel environment or social contact [61–63]. In fact, the induction of activity-dependent gene expression in the striatum and cortex is also observed after treatment with first-generation antipsychotics (e.g., haloperidol) at doses that cause catalepsy [64–66]. Nevertheless, these effects are due mainly to the activation of dopamine receptor D2-expressing neurons (i.e., the indirect pathway of medium spiny neurons in the striatum) [67]. We note, however, that unlike L-DOPA, the effects of drugs of abuse and some antipsychotics are particularly strong in the nucleus accumbens. Thus, while the induction of activity-regulated transcripts likely reflects a universal pattern linked to the activation of neurons, the spatial pattern differs depending on the drug.

Our results revealed several transcripts whose expression was affected by L-DOPA but was restricted to specific brain areas. Consistent with previous reports, we found that treatment induced the expression of *Nptx2,* which encodes neuronal pentraxin II (also known as Narp), a protein involved in the clustering of glutamate receptors, in the dorsal striatum [26]. Changes in *Nptx2* expression are linked to the development of L-DOPA-induced dyskinesias, and the presence of the protein product was reported in substantia nigra pars compacta Lewy bodies in a human postmortem PD study [68].

Here, we found that the drug-induced increase in *Nptx2* expression was specific to the dorsal striatum (cluster #0), with its transcript levels unaltered in other areas of the basal ganglia or dorsal cortex. Six additional transcripts also exhibited striatum-specific induction, namely, *Atg101* (autophagy-related protein 101), *Hspa4* (heat shock 70 kDa protein 4), *Vgf* (VGF nerve growth factor inducible), *Trak1* (trafficking kinesin protein 1), *Ist1* (IST1 factor associated with ESCRT-III) and *Adam23* (a disintegrin and metalloproteinase domain-containing protein 23). The first four of these transcripts share a similar pattern of expression to *Nptx2*, with relatively high levels in layers 2/3 of the dorsal cortex (cluster #3), low abundance in the nucleus accumbens (#2), and a significant L-DOPA-dependent increase in expression in the dorsal striatum (#0). The functional link between these transcripts remains unclear. Notably, the spatial methodology used here does not have single-cell resolution, and the possibility that the transcripts are induced in different cell types cannot be excluded.

A well-defined pattern of gene expression that was unique to the ventral cortex area corresponding to layer 1 of the piriform cortex (cluster #17) was also observed, which included the *Pqbp1* (polyglutamine binding protein-1), *Pnrc1* (proline-rich nuclear receptor coactivator 1), *Rmnd5a* (required for meiotic nuclear division 5 homolog A), and *Elmo1* (engulfment and cell motility protein 1) transcripts. The first three encode nuclear proteins with complex functions that form large multiprotein complexes and are involved in the processing or transcription of RNAs, whereas *Elmo1* has been reported to be involved in spine formation and neuronal plasticity. Moreover, the majority of significant L-DOPA-induced decreases in transcript abundance were also observed in the piriform cortex and the endopiriform nucleus. Examples include *Dtnb1* (dystrobrevin binding protein 1, #17), *Angel2* (Angel homolog 2, #21) and *Pnck* (calcium/calmodulin-dependent protein kinase type 1B, #14). The patterns of L-DOPA-induced decreases differed among the three areas, which is unsurprising considering that each is dominated by different neuronal populations, i.e., GABAergic interneurons, glutamatergic projection neurons, and multipolar GABAergic cells. Furthermore, the piriform cortex is linked primarily to the sense of smell; thus, changes induced by the degeneration of dopaminergic neurons and L-DOPA treatment are consistent with the impairments in olfactory acuity frequently observed in the early stages of PD. However, as we have previously shown, dopaminergic cells in the olfactory system of the brain of TIF-IA^DATCreERT2^ mice are not affected by the mutation (due to little to no expression of DAT). As shown here and in our previous reports, the dopaminergic projections crossing the olfactory tubercle are still present at the stage selected for analysis here [33,35]. Accordingly, we previously reported normal olfactory acuity in TIF-IA^DATCreERT2^ mice, although arguably, the task used was relatively simple, and a mild impairment could have had no effect. Thus, while L-DOPA-induced changes in gene expression in the piriform cortex could be interpreted as linked to an impaired sense of smell in PD patients, we would also like to point to a different aspect of its function. The piriform cortex is closely linked to the limbic system and has been shown to play a role in the attribution of valence to sensory stimuli, including visual stimuli, in humans [69–72]. Speculatively, this pathway could affect operant behavior and be linked to the increase in instrumental responding observed in TIF-IA^DATCreERT2^ mice (observed here and reported previously [35]). We do note, however, that even mild changes in the mesolimbic dopaminergic system that remain below the threshold needed to result in significant L-DOPA-induced changes in gene expression in the nucleus accumbens (clusters #2 and #11) would be the most likely link to the mechanism underlying this phenotype. Taken together, these observations reveal a complex effect of L-DOPA on brain areas involved in the nonmotor symptoms of PD.

Finally, the pattern of changes in gene expression in cluster #25 should also be highlighted, as it confirms the advantage of spatial analysis over other methods. The cluster has a scattered distribution in the dorsal striatum, which resembles the white matter tracts crossing the dorsal striatum (the eponymous stripes). The clustering analysis of gene expression indicated that this area is similar to other striatal areas (e.g., #0 and #2) rather than white matter (#10). Strikingly, unlike the surrounding dorsal striatum, L-DOPA caused predominant decreases in transcript abundance (e.g., *Parl*, presenilin-associated rhomboid-like protein), with only *Junb* and *Chga* (chromogranin A) expression significantly induced by treatment. While the identities of the cells in #25 remain elusive, they are clearly defined spatially and appear to be functionally different from those of the surrounding areas.

This study shows the feasibility of a spatial analysis of drug-induced changes in gene expression. We found that in a mouse model of progressive dopaminergic neuron degeneration, L-DOPA induced a spatially discrete pattern of gene expression, with the robust induction of activity-regulated gene expression in the dorsal striatum and the dorsal and ventral cortices. Furthermore, the analysis revealed different and, in some cases, unique drug-induced gene expression signatures that may be relevant to the nonmotor phenotypes associated with dopaminergic neuron degeneration.

## AUTHORS’ CONTRIBUTION

ARB, JP, MP and JRP designed the study; ARB performed all the behavioral experiments and animal procedures with ŁS and JRP; ARB and M Ziemiańska performed all the Visium molecular and histological procedures; SG, MC and JH performed the Nanopore study; ARB, M Zięba, MP, and JRP prepared the figures; ARB, MP, M Zięba, JH, MK and JRP analyzed the data; MB and GK generated and provided the genetically modified mice used in the study; and ARB and JRP wrote the manuscript with help from all the authors.

## DATA AVAILABILITY STATEMENT

RNA-seq data generated in the experiment is deposited in the Sequence Read Archive (https://www.ncbi.nlm.nih.gov/sra) under BioProject accession no. PRJNA1080215. Data regarding behavioral phenotyping, microscopic images and files used in spatial transcriptomic analyses are available at Zenodo webpage of the project: https://zenodo.org/records/14762864. Scripts used to perform transcriptomic (https://github.com/annaradli/tif-ldopa-slide1/tree/main) and behavioral analyses (https://github.com/annaradli/tif-ldopa-phenotype) can be found at their respective GitHub project sites.

## ETHICS APPROVAL

The experiments were planned and executed in accordance with the European and Polish laws concerning the use and welfare of laboratory animals (Directive 2010/63/UE, European Convention for the Protection of Vertebrate Animals Used for Experimental and other Scientific Purposes ETS No. 123, and Polish Law Dz.U. 2015 poz. 266). All procedures were approved by the II Local Ethical Committee in Krakow (permits no. 197/2021 and 92/2022).

## REFERENCES

1. Braak H, Ghebremedhin E, Rüb U, Bratzke H, Del Tredici K. Stages in the development of Parkinson’s disease-related pathology. Cell Tissue Res. 2004;318:121–134.

2. Hornykiewicz O. Biochemical aspects of Parkinson’s disease. Neurology. 1998;51:S2–S9.

3. Poewe W, Seppi K, Tanner CM, Halliday GM, Brundin P, Volkmann J, et al. Parkinson disease. Nat Rev Dis Primers. 2017;3:1–21.

4. Dauer W, Przedborski S. Parkinson’s Disease: Mechanisms and Models. Neuron. 2003;39:889–909.

5. Schapira AHV, Chaudhuri KR, Jenner P. Non-motor features of Parkinson disease. Nat Rev Neurosci. 2017;18:435–450.

6. Poewe W. Non-motor symptoms in Parkinson’s disease. European Journal of Neurology. 2008;15:14–20.

7. Kalia LV, Lang AE. Parkinson’s disease. The Lancet. 2015;386:896–912.

8. Lloyd KG, Davidson L, Hornykiewicz O. The neurochemistry of Parkinson’s disease: effect of L-dopa therapy. J Pharmacol Exp Ther. 1975;195:453–464.

9. Cotzias GC, Papavasiliou PS, Gellene R. Modification of Parkinsonism — Chronic Treatment with L-Dopa. New England Journal of Medicine. 1969;280:337–345.

10. Jenner P. Molecular mechanisms of L-DOPA-induced dyskinesia. Nat Rev Neurosci. 2008;9:665– 677.

11. Cenci MA. Presynaptic Mechanisms of l-DOPA-Induced Dyskinesia: The Findings, the Debate, and the Therapeutic Implications. Front Neurol. 2014;5.

12. Nagatsu T, Sawada M. L-dopa therapy for Parkinson’s disease: Past, present, and future. Parkinsonism & Related Disorders. 2009;15:S3–S8.

13. De Deurwaerdère P, Di Giovanni G, Millan MJ. Expanding the repertoire of L-DOPA’s actions: A comprehensive review of its functional neurochemistry. Progress in Neurobiology. 2017;151:57–100.

14. Skovgård K, Barrientos SA, Petersson P, Halje P, Cenci MA. Distinctive Effects of D1 and D2 Receptor Agonists on Cortico-Basal Ganglia Oscillations in a Rodent Model of L-DOPA-Induced Dyskinesia. Neurotherapeutics. 2023;20:304–324.

15. Cools R. Dopaminergic modulation of cognitive function-implications for l-DOPA treatment in Parkinson’s disease. Neuroscience & Biobehavioral Reviews. 2006;30:1–23.

16. Berke JD, Paletzki RF, Aronson GJ, Hyman SE, Gerfen CR. A Complex Program of Striatal Gene Expression Induced by Dopaminergic Stimulation. J Neurosci. 1998;18:5301–5310.

17. Morelli M, Cozzolino A, Pinna A, Fenu S, Carta A, Di Chiara G. l-Dopa stimulates c-*fos* expression in dopamine denervated striatum by combined activation of D-1 and D-2 receptors. Brain Research. 1993;623:334–336.

18. Svenningsson P, Gunne L, Andren PE. l-DOPA produces strong induction of c-fos messenger RNA in dopamine-denervated cortical and striatal areas of the common marmoset. Neuroscience. 2000;99:457–468.

19. Cole DG, Growdon JH, DiFiglia M. Levodopa Induction of Fos Immunoreactivity in Rat Brain Following Partial and Complete Lesions of the Substantia Nigra. Experimental Neurology. 1993;120:223–232.

20. Ebihara K, Ishida Y, Takeda R, Abe H, Matsuo H, Kawai K, et al. Differential expression of FosB, c-Fos, and Zif268 in forebrain regions after acute or chronic l-DOPA treatment in a rat model of Parkinson’s disease. Neuroscience Letters. 2011;496:90–94.

21. Carta AR, Tronci E, Pinna A, Morelli M. Different responsiveness of striatonigral and striatopallidal neurons to L-DOPA after a subchronic intermittent L-DOPA treatment. European Journal of Neuroscience. 2005;21:1196–1204.

22. Zhang X, Andren PE, Svenningsson P. Repeated l-DOPA treatment increases c-fos and BDNF mRNAs in the subthalamic nucleus in the 6-OHDA rat model of Parkinson’s disease. Brain Research. 2006;1095:207–210.

23. Ostock CY, Dupre KB, Eskow Jaunarajs KL, Walters H, George J, Krolewski D, et al. Role of the primary motor cortex in l-DOPA-induced dyskinesia and its modulation by 5-HT1A receptor stimulation. Neuropharmacology. 2011;61:753–760.

24. Atifi-Borel ME, Buggia-Prevot V, Platet N, Benabid A-L, Berger F, Sgambato-Faure V. De novo and long-term l-Dopa induce both common and distinct striatal gene profiles in the hemiparkinsonian rat. Neurobiology of Disease. 2009;34:340–350.

25. Han C-L, Liu Y-P, Sui Y-P, Chen N, Du T-T, Jiang Y, et al. Integrated transcriptome expression profiling reveals a novel lncRNA associated with L-DOPA-induced dyskinesia in a rat model of Parkinson’s disease. Aging (Albany NY). 2020;12:718–739.

26. Charbonnier-Beaupel F, Malerbi M, Alcacer C, Tahiri K, Carpentier W, Wang C, et al. Gene Expression Analyses Identify Narp Contribution in the Development of l-DOPA-Induced Dyskinesia. J Neurosci. 2015;35:96–111.

27. Konradi C, Westin JE, Carta M, Eaton ME, Kuter K, Dekundy A, et al. Transcriptome analysis in a rat model of l-DOPA-induced dyskinesia. Neurobiology of Disease. 2004;17:219–236.

28. Smith LM, Parr-Brownlie LC, Duncan EJ, Black MA, Gemmell NJ, Dearden PK, et al. Striatal mRNA expression patterns underlying peak dose l-DOPA-induced dyskinesia in the 6-OHDA hemiparkinsonian rat. Neuroscience. 2016;324:238–251.

29. Heiman M, Heilbut A, Francardo V, Kulicke R, Fenster RJ, Kolaczyk ED, et al. Molecular adaptations of striatal spiny projection neurons during levodopa-induced dyskinesia. Proc Natl Acad Sci U S A. 2014;111:4578–4583.

30. Klemann CJHM, Xicoy H, Poelmans G, Bloem BR, Martens GJM, Visser JE. Physical Exercise Modulates L-DOPA-Regulated Molecular Pathways in the MPTP Mouse Model of Parkinson’s Disease. Mol Neurobiol. 2018;55:5639–5657.

31. Figge DA, Amaral H de O, Crim J, Cowell RM, Standaert DG, Jaunarajs KLE. Differential Activation States of Direct Pathway Striatal Output Neurons during l-DOPA-Induced Dyskinesia Development. J Neurosci. 2024;44.

32. Radlicka A, Kamińska K, Borczyk M, Piechota M, Korostyński M, Pera J, et al. Effects of L-DOPA on Gene Expression in the Frontal Cortex of Rats with Unilateral Lesions of Midbrain Dopaminergic Neurons. eNeuro. 2021;8.

33. Rieker C, Engblom D, Kreiner G, Domanskyi A, Schober A, Stotz S, et al. Nucleolar disruption in dopaminergic neurons leads to oxidative damage and parkinsonism through repression of mammalian target of rapamycin signaling. J Neurosci. 2011;31:453–460.

34. Kreiner G, Rafa-Zabłocka K, Barut J, Chmielarz P, Kot M, Bagińska M, et al. Stimulation of noradrenergic transmission by reboxetine is beneficial for a mouse model of progressive parkinsonism. Sci Rep. 2019;9:5262.

35. Radlicka-Borysewska A, Jabłońska J, Lenarczyk M, Szumiec Ł, Harda Z, Bagińska M, et al. Non-motor symptoms associated with progressive loss of dopaminergic neurons in a mouse model of Parkinson’s disease. Front Neurosci. 2024;18.

36. Sert NP du, Ahluwalia A, Alam S, Avey MT, Baker M, Browne WJ, et al. Reporting animal research: Explanation and elaboration for the ARRIVE guidelines 2.0. PLOS Biology. 2020;18:e3000411.

37. Jastrzębska K, Walczak M, Cieślak PE, Szumiec Ł, Turbasa M, Engblom D, et al. Loss of NMDA receptors in dopamine neurons leads to the development of affective disorder-like symptoms in mice. Sci Rep. 2016;6:37171.

38. Olsen CM, Winder DG. Operant sensation seeking engages similar neural substrates to operant drug seeking in C57 mice. Neuropsychopharmacology. 2009;34:1685–1694.

39. Parkitna JR, Sikora M, Gołda S, Gołembiowska K, Bystrowska B, Engblom D, et al. Novelty-seeking behaviors and the escalation of alcohol drinking after abstinence in mice are controlled by metabotropic glutamate receptor 5 on neurons expressing dopamine d1 receptors. Biol Psychiatry. 2013;73:263–270.

40. Antunes M, Biala G. The novel object recognition memory: neurobiology, test procedure, and its modifications. Cogn Process. 2012;13:93–110.

41. Leger M, Quiedeville A, Bouet V, Haelewyn B, Boulouard M, Schumann-Bard P, et al. Object recognition test in mice. Nat Protoc. 2013;8:2531–2537.

42. Friard O, Gamba M. BORIS: a free, versatile open-source event-logging software for video/audio coding and live observations. Methods in Ecology and Evolution. 2016;7:1325–1330.

43. Simon P, Dupuis R, Costentin J. Thigmotaxis as an index of anxiety in mice. Influence of dopaminergic transmissions. Behavioural Brain Research. 1994;61:59–64.

44. Gould TD, Dao DT, Kovacsics CE. The Open Field Test. In: Gould TD, editor. Mood and Anxiety Related Phenotypes in Mice: Characterization Using Behavioral Tests, Totowa, NJ: Humana Press; 2009. p. 1–20.

45. Paxinos G, Franklin KBJ. The Mouse Brain in Stereotaxic Coordinates (Deluxe Edition), Second Edition. 2nd edition. San Diego: Academic Press; 2001.

46. Mustafa R, Rawas C, Mannal N, Kreiner G, Spittau B, Kamińska K, et al. Targeted Ablation of Primary Cilia in Differentiated Dopaminergic Neurons Reduces Striatal Dopamine and Responsiveness to Metabolic Stress. Antioxidants (Basel). 2021;10:1284.

47. Li H, Handsaker B, Wysoker A, Fennell T, Ruan J, Homer N, et al. The Sequence Alignment/Map format and SAMtools. Bioinformatics. 2009;25:2078–2079.

48. Chrószcz M, Hajto J, Misiołek K, Szumiec Ł, Ziemiańska M, Radlicka-Borysewska A, et al. μ-Opioid receptor transcriptional variants in the murine forebrain and spinal cord. Gene. 2025;932:148890.

49. Stuart T, Butler A, Hoffman P, Hafemeister C, Papalexi E, Mauck WM, et al. Comprehensive Integration of Single-Cell Data. Cell. 2019;177:1888–1902.e21.

50. 50. Reference Atlas :: Allen Brain Atlas: Mouse Brain. https://mouse.brain-map.org/static/atlas. Accessed 26 January 2025.

51. Gerfen CR, Scott Young W. Distribution of striatonigral and striatopallidal peptidergic neurons in both patch and matrix compartments: an in situ hybridization histochemistry and fluorescent retrograde tracing study. Brain Research. 1988;460:161–167.

52. Gokce O, Stanley GM, Treutlein B, Neff NF, Camp JG, Malenka RC, et al. Cellular Taxonomy of the Mouse Striatum as Revealed by Single-Cell RNA-Seq. Cell Rep. 2016;16:1126–1137.

53. Muñoz-Castañeda R, Zingg B, Matho KS, Chen X, Wang Q, Foster NN, et al. Cellular anatomy of the mouse primary motor cortex. Nature. 2021;598:159–166.

54. Smith JB, Alloway KD, Hof PR, Orman R, Reser DH, Watakabe A, et al. The relationship between the claustrum and endopiriform nucleus: a perspective towards consensus on cross-species homology. J Comp Neurol. 2019;527:476–499.

55. Piñero J, Ramírez-Anguita JM, Saüch-Pitarch J, Ronzano F, Centeno E, Sanz F, et al. The DisGeNET knowledge platform for disease genomics: 2019 update. Nucleic Acids Research. 2020;48:D845–D855.

56. Graybiel AM, Moratalla R, Robertson HA. Amphetamine and cocaine induce drug-specific activation of the c-fos gene in striosome-matrix compartments and limbic subdivisions of the striatum. Proc Natl Acad Sci USA. 1990;87:6912–6916.

57. Piechota M, Korostynski M, Solecki W, Gieryk A, Slezak M, Bilecki W, et al. The dissection of transcriptional modules regulated by various drugs of abuse in the mouse striatum. Genome Biol. 2010;11:R48.

58. Piechota M, Golda S, Ficek J, Jantas D, Przewlocki R, Korostynski M. Regulation of alternative gene transcription in the striatum in response to antidepressant drugs. Neuropharmacology. 2015;99:328–336.

59. Korostynski M, Piechota M, Dzbek J, Mlynarski W, Szklarczyk K, Ziolkowska B, et al. Novel drug-regulated transcriptional networks in brain reveal pharmacological properties of psychotropic drugs. BMC Genomics. 2013;14:606.

60. Zygmunt M, Piechota M, Rodriguez Parkitna J, Korostyński M. Decoding the transcriptional programs activated by psychotropic drugs in the brain. Genes Brain Behav. 2018:e12511.

61. Kabbaj M, Akil H. Individual differences in novelty-seeking behavior in rats: a c-*fos* study. Neuroscience. 2001;106:535–545.

62. Rinaldi A, Romeo S, Agustín-Pavón C, Oliverio A, Mele A. Distinct patterns of Fos immunoreactivity in striatum and hippocampus induced by different kinds of novelty in mice. Neurobiology of Learning and Memory. 2010;94:373–381.

63. Ahern M, Goodell DJ, Adams J, Bland ST. Brain regional differences in social encounter-induced Fos expression in male and female rats after post-weaning social isolation. Brain Research. 2016;1630:120–133.

64. Robertson GS, Fibiger HC. Neuroleptics increase c-fos expression in the forebrain: contrasting effects of haloperidol and clozapine. Neuroscience. 1992;46:315–328.

65. Ziółkowska B, Höllt V. Fos is not involved in the regulation of the proenkephalin gene by haloperidol in the mouse striatum. Molecular Brain Research. 1995;34:351–354.

66. Deutch AY, Duman RS. The effects of antipsychotic drugs on Fos protein expression in the prefrontal cortex: cellular localization and pharmacological characterization. Neuroscience. 1996;70:377–389.

67. Bertran-Gonzalez J, Bosch C, Maroteaux M, Matamales M, Hervé D, Valjent E, et al. Opposing patterns of signaling activation in dopamine D1 and D2 receptor-expressing striatal neurons in response to cocaine and haloperidol. J Neurosci. 2008;28:5671–5685.

68. Moran LB, Hickey L, Michael GJ, Derkacs M, Christian LM, Kalaitzakis ME, et al. Neuronal pentraxin II is highly upregulated in Parkinson’s disease and a novel component of Lewy bodies. Acta Neuropathol. 2008;115:471–478.

69. Blazing RM, Franks KM. Odor coding in piriform cortex: mechanistic insights into distributed coding. Current Opinion in Neurobiology. 2020;64:96–102.

70. Nordén F, Iravani B, Schaefer M, Winter AL, Lundqvist M, Arshamian A, et al. The human olfactory bulb communicates perceived odor valence to the piriform cortex in the gamma band and receives a refined representation back in the beta band. PLoS Biol. 2024;22:e3002849.

71. Schulze P, Bestgen A-K, Lech RK, Kuchinke L, Suchan B. Preprocessing of emotional visual information in the human piriform cortex. Sci Rep. 2017;7:9191.

72. Bekkers JM, Suzuki N. Neurons and circuits for odor processing in the piriform cortex. Trends in Neurosciences. 2013;36:429–438.

